# Automated leg tracking reveals distinct conserved gait and tremor signatures in *Drosophila* models of Parkinson’s Disease and Spinocerebellar ataxia 3

**DOI:** 10.1101/425405

**Authors:** Shuang Wu, Kah Junn Tan, Lakshmi Narasimhan Govindarajan, James Charles Stewart, Lin Gu, Joses Wei Hao Ho, Malvika Katarya, Boon Hui Wong, Eng-King Tan, Daiqin Li, Adam Claridge-Chang, Camilo Libedinsky, Li Cheng, Sherry Shiying Aw

## Abstract

Genetic models in *Drosophila* have made invaluable contributions to our understanding of the molecular mechanisms underlying neurodegeneration. In human patients, some neurodegenerative diseases lead to characteristic movement dysfunctions, such as abnormal gait and tremors. However, it is currently unknown whether similar movement defects occur in the respective fly models, which could be used to model and better understand the pathophysiology of movement disorders. To address this question, we developed a machine-learning image-analysis programme — Feature Learning-based LImb segmentation and Tracking (FLLIT) — that automatically tracks leg claw positions of freely moving flies recorded on high-speed video, generating a series of body and leg movement parameters. Of note, FLLIT requires no user input for learning. We used FLLIT to characterise fly models of Parkinson’s Disease (PD) and Spinocerebellar ataxia 3 (SCA3). Between these models, walking gait and tremor characteristics differed markedly, and recapitulated signatures of the respective human diseases. Selective expression of mutant SCA3 in dopaminergic neurons led to phenotypes resembling that of PD flies, suggesting that the behavioural phenotype may depend on the circuits affected, rather than the specific nature of the mutation. Different mutations produced tremors in distinct leg pairs, indicating that different motor circuits are affected. Almost 190,000 video frames were tracked in this study, allowing, for the first time, high-throughput analysis of gait and tremor features in *Drosophila* mutants. As an efficient assay of mutant gait and tremor features in an important model system, FLLIT will enable the analysis of the neurogenetic mechanisms that underlie movement disorders.

## Introduction

Walking requires coordination of the central and peripheral nervous systems, and the musculoskeletal system. Hence, neurological and musculoskeletal pathologies can manifest as movement abnormalities. For example, patients with Parkinson’s disease (PD) exhibit slowed movements (bradykinesia), rigidity and resting tremor^1^, while patients with cerebellar ataxias like Spinocerebellar ataxia Type 3 (SCA3) exhibit stumbling, jerky, uncoordinated movements, and action tremor^2-5^. The gait and tremor movements exhibited by patients give clues as to the affected brain regions, and are used to inform diagnosis^6^. Pathological gait and tremor cause difficulty for basic tasks required for daily living; yet, the mechanisms and affected neuronal circuitry are poorly described. One common disease signature is tremor: uncontrolled shaking of the body or appendages. While tremors can be temporarily triggered by physiological states like stress, pathological tremors are often symptomatic of an underlying neurological disorder. There is no cure for tremor, and its pathophysiological causes are not understood^7,8^. Movement disorders exhibit significant phenotypic heterogeneity^3,9-11^; hence, detailed characterisation of circuit dysfunction can aid in understanding the factors that influence disease severity and progression. To achieve this level of understanding requires linking detailed behavioural measurements with functional circuit analyses.

Despite the differences in anatomy and scale between human and fly brains, *Drosophila* disease models have made substantial contributions to our understanding of the mechanisms underlying human neurodegenerative diseases^12,13^. However, while fly disease models recapitulate molecular features of disease, and fly and mammalian neurons share similar molecular machinery^14^, it is unknown whether flies can be used to model movement disorders, as gait and tremor characterisatics of fly neurodegeneration models have not been quantified. Conserved cellular roles for human disease genes may not result in conserved movement dysfunctions, which depend on circuit properties. Tremor behaviour, in particular, has eluded characterisation; while flies of several mutant genotypes have been reported to exhibit trembling behaviour^15-18^, the quantitative characteristics of these tremors have not been determined.

To characterise movement defects and tremors in flies requires an accurate, automated method for fly leg tracking. State-of-the-art methods include foot-printing-based approaches that report contact points with a detection surface^19,20^, and leg marking-based techniques that track distinct marked spots on the legs^21^. Semi-automated algorithms have been developed to aid high-resolution leg tracking in freely-moving, unmarked flies^22-24^, but these require a considerable degree of user annotation and/or user-led optimisation. Therefore, these methods were not feasible for use on the large volume of data required to quantify rapid and fine tremors in suspended legs. Two recent studies describe deep learning approaches for markerless tracking^25,26^. These methods require a substantial number of user annotated images for training, however, are versatile, as they can be applied to various models and behaviours.

To enable accurate, automated leg tracking, we developed FLLIT (Feature Learning-based LImb segmentation and Tracking), a machine learning method able to automatically track leg movements of freely moving flies from high-speed video, with high accuracy and minimal user input. Importantly, FLLIT does not require user annotation of images for training, which sets it apart from other learning approaches^25,26^. Using FLLIT to characterise gait in fly models for PD and SCA3, we found that these models exhibit distinct movement signatures that recapitulate aspects of the movement dysfunctions in human patients. Such gait and tremor analyses can enable future studies into the genetic and neural basis underlying subtle limb behaviours and movement defects like tremors.

## Results

### 1. System setup and computational workflow

We utilised a video recording setup that consists of a high-speed camera mounted below the sample, backlit by an infrared LED array (Figs. 1A, S1). The computational workflow carried out by FLLIT consists of: (i) Automated training set generation, (ii) Supervised learning of leg classification, (iii) Application of the trained classifier to novel images for leg segmentation, (iv) Leg tracking across frames, and (v) Results output (Fig. 1B). A brief overview of each stage of the workflow is detailed below.

### Generating the training set

To identify and segment legs in each video frame, we use an iterative learning approach that incorporates steps for the automated preparation of training sets. This circumvents the normally tedious manual preparation of training samples, as no user input is required for training. To achieve this, we perform a series of image processing steps (Fig. 1Bi) on a subset of the images from the video of interest: background subtraction^27^, medial axis skeletonisation and edge extraction, to identify only high-confidence positive (leg) and negative (non-leg) pixel examples for learning.

### Identification of legs

Our iterative learning approach is based on the KernelBoost method^28,29^, and aims to learn a precise binary leg segmentation model from pixels in the training set (Fig. 1Bii). Training examples are extracted as square image patches consisting of a central pixel of interest and surrounding pixels. Candidate filters (convolution kernels) are applied to confer features onto each image patch, which are then used to learn parameters on decision trees (weak learners). High-confidence pixel classification predictions at the current iteration are added to the training set of the next iteration, and thus iteratively augment the training set in favour of the safe and/or high-confidence predictions. The feedback and retraining loops expand the training set to build a stronger segmentation model at each iteration (Fig. 1Bii). After the classifier has been trained, it can be used to predict leg pixels in novel images (based on a set classification threshold) (Fig. 1Biii), and the predicted leg pixels are grouped as legs. These segmented leg images are saved by FLLIT, and can be used for further analysis.

### Leg tracking

After the legs have been identified, tracking can be performed (Fig. 1Biv and Video 1). Each frame is body centred, and leg claw positions are matched across adjacent frames using the Hungarian method^30^. The tracked leg-tip data for each leg can then be extracted as a CSV file for analysis (Fig. 1Bv).

**Figure 1.**
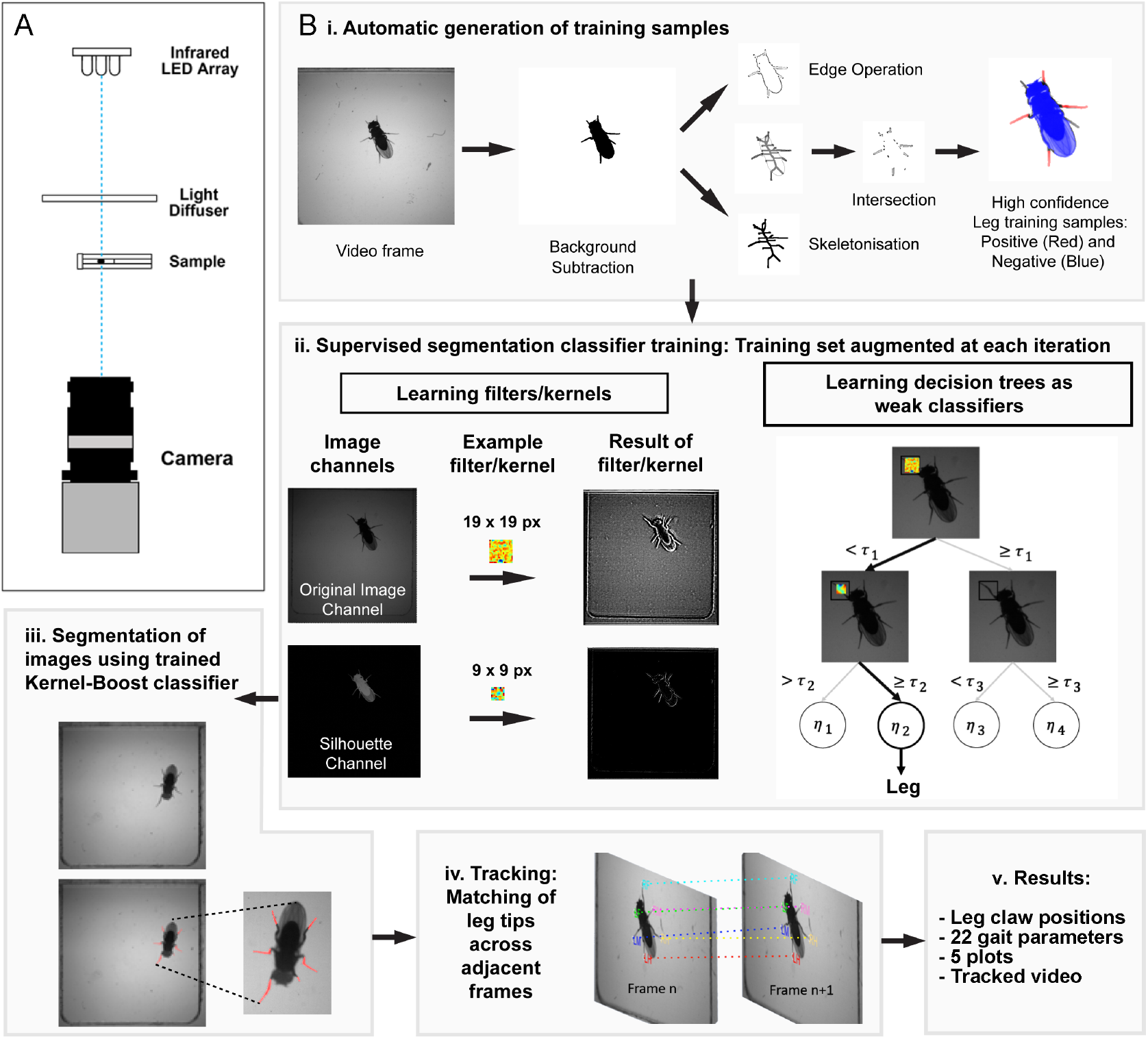
FLLIT System setup and overview of computational workflow. **A.** Camera and arena setup used for video capture. **B.** Segmentation and tracking procedure. i) Training samples are automatically generated by identifying high confidence leg pixels (px) located at the intersection between skeletonisation and edge morphological operations. ii) Training sets are learned and grown by iterative supervised segmentation to derive a classifier. iii) Segmentation of novel images is carried out using the trained classifier. iv) Tracking occurs by matching leg claw positions across adjacent frames. v) Results are given as positions of leg claws in each frame.

### FLLIT software

The FLLIT software provides an interface for automated leg tracking of high-speed videos. Besides the set of data files of tracked body and leg positions, it also automatically calculates 23 body and gait parameters, and provides 5 plots and a video for visualising the tracked data (Table 1). The FLLIT program, readme and sample data can be downloaded from: https://github.com/BII-wushuang/FLLIT

**Table 1.** Movement and gait parameters produced by FLLIT. include raw body and leg claw position data, as well as 23 gait parameters, 5 plots and a tracked video.

| FLLIT Parameters |  |
| --- | --- |
| Raw data |  |
|  | Body position |
|  | Leg claw positions |
| Body parameters |  |
|  | Body trajectory |
|  | Body length |
|  | Instantaneous body velocity |
|  | Turning points of the body trajectory |
| Stride parameters (per leg) |  |
|  | Stride duration |
|  | Stride period |
|  | Stride displacement/length |
|  | Stride path covered |
|  | Take-off position |
|  | Landing position |
|  | Stride amplitude |
|  | Stance linearity |
|  | Stance width |
| Leg parameters (per leg) |  |
|  | Instantaneous speed of each leg |
|  | Gait index |
|  | Percentage of time in motion |
|  | Mean stride period |
|  | Footprint regularity |
|  | Leg trajectory domain area |
|  | Leg trajectory domain length and width |
|  | Leg trajectory domain intersection |
|  | Footprint deviation from trajectory principal component |
|  | Foot-dragging event count |
| Plots/Video |  |
|  | Body trajectory |
|  | Body velocity |
|  | Leg trajectories in body-centred frame of reference |
|  | Gait plots using speed |
|  | Gait index plot |
|  | Video of tracked data in arena- and body-centred views |

## 2. Accuracy of FLLIT segmentation and tracking results determined by ground truth

To determine an optimal classification confidence threshold, we manually identified leg pixels in video frames of wild-type flies, and examined classifier performance at different thresholds (Fig. S2A). A threshold of 0.65 was selected for subsequent tracking analyses, based on the F_0.5_ scores for the tested classifiers, which peaked at 0.6–0.65. We then interrogated the level of similarity that can be expected between leg-tip annotations of the same set of images made by two individuals. Pixel deviation was measured as the Euclidean distance between the pixels selected by the two individuals (Fig. 2A). We found that different individuals asked to identify leg claw positions in the same set of images located the same pixels (0 pixel deviation) about 33% of the time (Fig. 2B). The majority of tips were located 1–2 pixels apart. At our recording resolution, this deviation corresponded to ~ 0.75-1.5% of body length, which averaged 133 pixels (Fig. S2B).

We then compared the leg-tip positions reported by the algorithm to those determined by manual annotation. We found three categories of errors: 1) Misidentification errors; 2) Missing data of occluded or visible legs; 3) Deviation errors. These errors are detailed below.

## Misidentification errors

Misidentification errors occurred when leg identities were erroneously assigned to the wrong body part, which can lead to tracking errors in subsequent frames. For example, in some strides, the forelegs may become occluded during leg retraction. Occluded objects cannot be detected using silhouette imaging. When trying to locate the occluded leg claw, the algorithm may falsely label the antenna or another leg (Fig. S2C). These errors are salient, and their correction can prevent perpetuation of the error through the rest of the video. We therefore corrected these errors before examining the rest of the tracked data (Fig. S2C). Absent leg claw positions that resulted from corrections were considered as missing data (next section). By this approach, we found that wild-type flies required an average of ~1.6 corrections for misidentifications per 1,000 frames; 72% of videos required no corrections for misidentifications (Fig. 2C). We used FLLIT-automated background subtraction for all datasets examined in this study, and noticed that inefficient background subtraction—which causes poor segmentation— contributed to misidentification errors. In these cases, loading of a background image, instead of using automated background subtraction, can improve segmentation and tracking (Fig. S2D).

## Missing values

Missing leg claw data occurred as a result of occlusion or failure to track visible legs. An average of ~3.6% of tips per video were not tracked (Fig. 2D). The majority (~86%) of missing values occurred in the front legs, which are partially (28%; 235/833 missing tips; e.g. Fig. S2E) or completely (72%; 598/833 missing tips, e.g. Fig. S2Ci) obscured by the body when they retract during each stride. Occasionally, hind legs were obscured by the wings. Overall, ~8.1% leg-tip data for the front legs were unreported, compared to 0.01% and 2.9% for the mid and hind legs, respectively (Fig. 2D).

## Deviation errors

Deviation errors occurred when the predicted leg-tip positions differed from the ground-truth annotated positions, and were determined after correction of misidentification errors. We found that almost 98% of reported tip locations were accurate to within 3 pixels (Fig. 2E). Of note, deviations of up to 3 pixels occurred by manual annotation (Fig. 2B), and leg tips spanned ~3 pixels in width under our recording parameters (Fig. 2A).

## Effect of learning on segmentation and tracking accuracy

We quantified the effect of learning on leg segmentation and tip tracking performance, compared to solely using morphological operations (as was used to derive the training set, Fig. 1Bi). As we selected only pixels of high confidence/precision for the initial training set (Fig. 1Bi), before learning, recall of leg pixels was low, while precision was high (Fig. 2F). After learning, recall scores showed stark improvement, demonstrating that the algorithm is able to generalize and classify additional leg pixels based on the narrow training set (Figs. 2F, S2F). This increase in recall does not result in decreased precision (Fig. 2F). We then examined tracking peformance after learning. The percentage of missing data decreased by ~31.5% on average for each sample (Fig. S2G). Of the found tips, deviation errors also decreased slightly (Fig. S2H).

In summary, our learning-based method can accurately track the leg-tip positions of freely walking, unmarked wild-type flies from high-speed video, and is more accurate than using morphological parameters alone.

**Figure 2.**
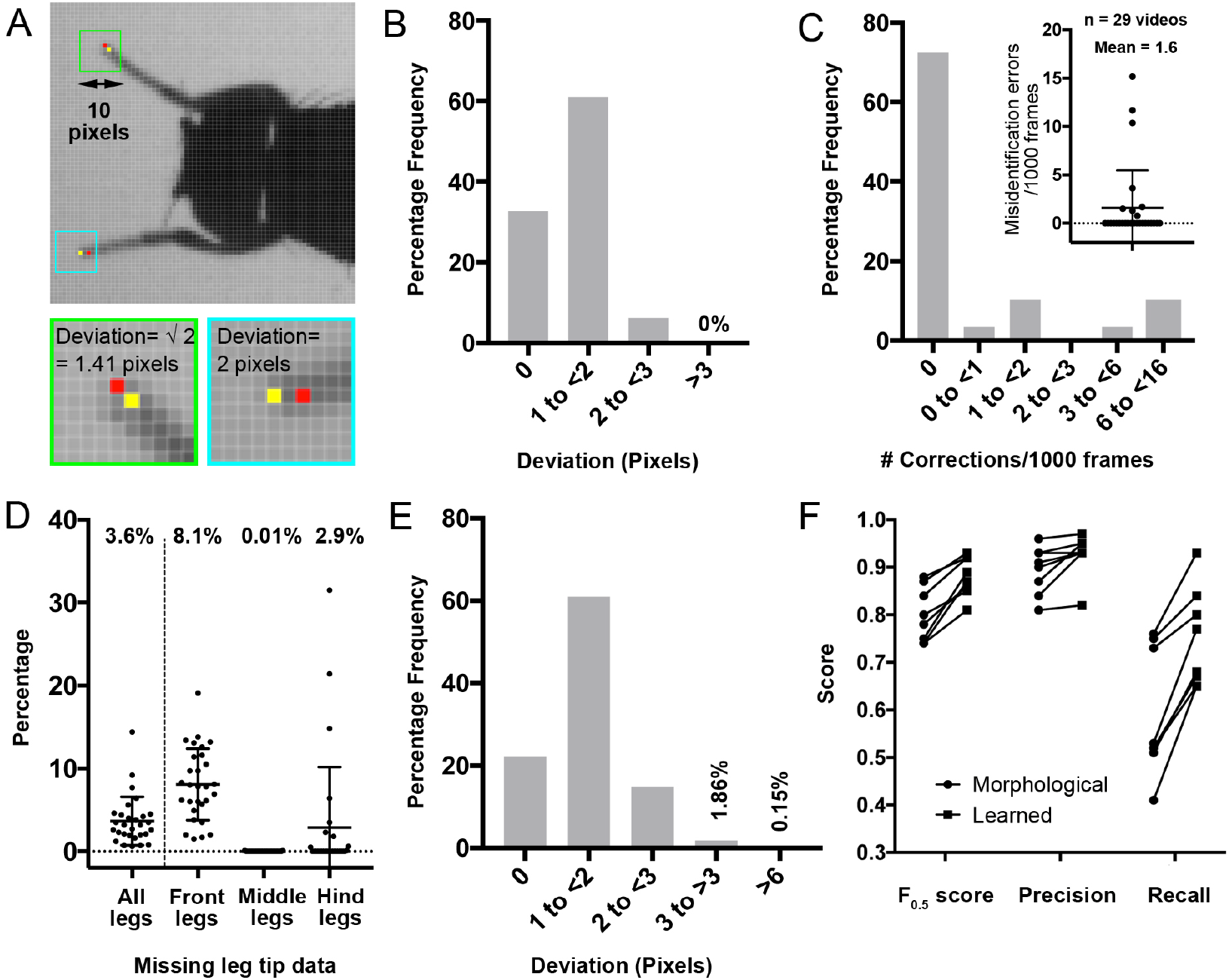
Ground truth demonstrates the accuracy of FLLIT (Feature Learning-based LImb segmentation and Tracking) segmentation and tracking results. **A.** Representative images of wild-type *Drosophila* legs taken using the default settings, and the manual leg-tip positions identified by two different human users. Blue and green insets are 10 pixels wide and show the respective boxed regions in the top image. Red and yellow dots represent the pixels identified as tip pixels by the two users, within the respective blue and green boxes. **B.** Frequency distribution of the deviation (in pixels) between leg-tip positions annotated by the two users (n = 54 frames, 324 leg tips, from two videos). Discrepancies can occur in both the X and Y directions, and are represented as the Euclidean distance between the two pixels. **C.** Number of corrections required for misidentified legs, normalised to per 1,000 frames (Mean = 1.7 corrections; n = 29 videos, 15,166 frames). Plotted as a frequency distribution and a scatter plot (inset). **D.** Percentage of missing data in wild-type Drosophila after tracking (n= 29 videos, 15,166 frames). **E.** Frequency distribution of the deviation (in pixels) between computationally and manually derived leg tip positions (n= 106 frames, 636 leg tips from two videos). **F.** Segmentation F_0.5_, precision and recall scores for each video, using only morphological parameters alone, or after learning and application of a FLLIT leg classifier. (n = 8 videos, 2-3 images per video). Bars represent the means and standard deviations. *See also Video 1*.

## 3. FLLIT robustness assessment

Image analysis software should be tolerant of deviations in recording parameters, as it is challenging to identically replicate a video recording setup from one lab to another. We examined the effect of altering the following parameters on FLLIT performance: 1) Image contrast — Low, default and high infrared light intensity; 2) Resolution — 9, 10 and 12 mm field of view; and 3) Video capture speed — 250, 500 and 1000 frames per second (fps) (Fig. S3A).

Upon altering video contrast, resolution and capture speed, the percentage of missing values ranged from 2.1–5.6% (Fig. S3Bi). The number of corrections needed to re-label misidentified legs ranged from ~0–9.3 per 1,000 frames (Fig. S3Bii). We then analyzed the final reported leg-tip positions. In all conditions, ~97.5% of the computationally identified tips were within 3 pixels of the manually annotated positions (Fig. S3Biii). These data support the conclusion that FLLIT can generate classifiers to track and analyse videos recorded under a range of settings, and is not dependent on a stringent set of conditions.

As FLLIT is not rule-based, we asked whether it could also be used to automatically track leg movements in other arthropods. As a test, we chose the *Myrmaplata plataleoides* salticid spider, which has eight legs and measures ~ 13 mm in length, and thus differs markedly from *Drosophila* in body plan and proportions (Fig. S3C). Salticid leg misidentifications occurred when legs touched or crossed over (mean = 1.2 corrections/1000 frames; n = 9 videos, 12,683 frames, 101,464 legs) (Fig. S3B). These corrections resulted in an average of ~0.66% of missing leg data (Fig. S3C). Computationally-predicted leg-tip positions compared favourably with manual annotation; >99% of the tracked data deviated by < 3 pixels from user-annotated positions (Fig. S3D). A small fraction deviated by >6 pixels, which was due to legs touching or crossing over during walking. In summary, these data suggest that FLLIT can be used to accurately track leg tips in other arthropods (Video S1).

## 4. Side-by-side performance comparison

Two types of approaches are currently available for automated *Drosophila* leg movement tracking (i.e. not requiring user input). One uses thresholding and dynamic masking methods (TDM) to automatically identify leg tips^22,31^, while two recent studies employ deep learning algorithms^25,26^ using user-annotated training sets. We first compared the performance of FLLIT against TDM-based software provided by Isakov, *et. al*^22^. We found that the TDM method was sensitive to image contrast, and required flies to walk sufficiently close to the centre of the video for thresholding. Hence, videos had to be cropped by trial and error for successful tracking. We directly compared the tracked positions, without making any error corrections for either method. The percentage of missing values when using either method was comparable (Fig. S4A). However, 16.1% of positions identified using TDM were located >3 pixels from the manually-derived positions, compared to 1.2% for FLLIT (>13 fold difference) (Fig. S4B). FLLIT also performed markedly better than TDM under other recording settings (Fig. S4C). Of note, we could not find parameters that allowed low contrast images to be tracked using TDM; these were accurately tracked by FLLIT (Figs. 3, S4A, C). To gauge the usefulness of a tracking tool, the magnitude of the deviation errors may matter less than the rate of error correction required. Hence, we assessed what percentage of frames required user correction (deviations >3 pixels; See text for Fig. 2). By the TDM method, ~32–63% of frames contained at least 1 one leg that deviated >3 pixels from the manually-derived positions, compared to 2.8–8.1% when using FLLIT (a 4-22 fold difference)(Fig. S4D).

A deep learning method for movement tracking, DeepLabCut, was recently published^26^. While FLLIT and the TDM method require no user input, DeepLabCut requires the user to pretrain the algorithm using at least 200 user-annotated images. We compared the performance of DeepLabCut to FLLIT in our leg tracking task. When DeepLabCut on was trained on a subset of images from a video, and used to test on novel images from the same video, 28.1% of frames predicted using DeepLabCut contained at least 1 leg deviating >3 pixels, compared to 6.3% with FLLIT (Fig. S4D). When DeepLab Cut was trained on images from one video, and then used to track a different video taken under similar settings, 100% of frames contained at least 1 leg deviating >3 pixels, even when we manually matched predicted leg claw positions to the closest leg (i.e. using DeepLabCut to find leg claw positions, without requiring labelling of leg identity). We therefore conclude that FLLIT performs substantially better on this task than currently available methods.

## 5. Characterisation of gait in *Drosophila* models of Spinocerebellar ataxia 3 (SCA3) and Parkinson’s Disease (PD)

Ataxic gait in SCA3 is typified by body veering, erratic foot placement, leg crossing over, lurching steps and intention/action tremor^5,32^, while gait in PD patients is marked by rigid, shuffling steps and resting tremor^1,32^ (Table 2). Therefore, these two diseases exhibit distinctly different gaits that arise from their underlying etiologies. We used FLLIT to determine gait characteristics of previously described fly models of the two diseases. For SCA3, we expressed wildtype and mutant human SCA3 under control of the pan-neuronal driver Elav-Gal4^33,34^. For PD we looked at two models: Expression of human alpha-synuclein (SCNA) under control of Elav-Gal4^35,36^, and *parkin* mutant flies^37-41^. These models exhibit neurodegeneration and gross motor defects reflected in poor climbing ability^33-41^. To enable comparison of gait defects amongst the different genotypes, we used climbing ability as a readout of phenotypic severity, selecting for analysis mutant flies with similar climbing performance (Fig. 3A).

**Table 2.** Gait features of Parkinson’s Disease and Spinocrebellar ataxia in human patients, and corresponding gait parameters used to analyse these features in FLLIT.

|  | Gait features of disease |  |  |  |  |
| --- | --- | --- | --- | --- | --- |
| Spinocerebellar ataxia 3 (SCA3) | Veering | Erratic foot placement and leg crossing over | Lurching steps | Short strides | Action tremor |
| Parkinson's Disease (PD) | Not a feature | Not a feature | Leg rigidity | Rigidity, Shuffling | Resting tremor |
| Measurement Parameter | Number of body turn events | Footprint regularity | Size of leg domains, degree of domain overlap | Stride length | Irregularities in leg traces |

Leg tracking with FLLIT showed that flies expressing wildtype human SCA3-flQ27 in all neurons walk coordinatedly, with strides that form regular leg domains (Fig. 3Bi, Video 2). However, pan-neuronal expression of SCA3-flQ84 mutants led to a strikingly aberrant gait (Fig. 3Bii, Video 3). Similar to that seen in SCA3 patients^5,32^, SCA3 flies exhibited repeated veering, detected as body turns (Figs. 3Ci, S5Ai). These uncoordinated movements and erratic foot placement resulted in low footprint regularity^19^, reflected in large standard deviations of the both the anterior (AEP, Figs. 3Ci, S5Aii) and posterior (not shown) extreme positions of the mid and hind legs, and longer (Figs. 3Ci, S5Aiii) leg domains of both the mid and hind legs. These leg domains were so large as to intersect with one another (Figs. 3Bii, 3Ci, S5Aiv).

While these phenotypes were striking in their similarity to cerebellar ataxic gait^42^, they could result from non-specific toxicity due to pan-neuronal expression of a pathogenic protein that is not endogenously expressed in *Drosophila*. We therefore examined gait in flies expressing alpha-synuclein (SCNA), the hallmark protein that accumulates in Lewy bodies in PD, which is also not endogenously present in *Drosophila*. Elav-Gal4-mediated expression of wild-type alpha-synuclein was previously shown to cause neurodegeneration and climbing defects^36^. We used a recently published codon-optimised version of UAS-alpha-synuclein with high Gal4-driven expression^35^. If both SCNA and SCA3 expression cause non-specific toxicity effects, flies of each genotype with similar climbing performance (Fig. 3A) should exhibit similar gait profiles. However, alpha-synuclein-expressing flies did not exhibit hyperkinetic movements of increased body turning, low footprint regularity, enlarged leg movement domains nor aberrant domain overlap (Figs. 3Biii, 3Cii, S5Bi-v, Video 4). To the contrary, they showed hypokinetic movements: leg rigidity in the form of short strides and *smaller* domain length especially in the hind legs, relative to the mid legs (Figs. 3Biii, 3Cii, S5Biii, iv, vi and vii, Video 4). Therefore, interestingly, expression of two different non-endogenous human disease proteins in all fly neurons not only result in distinct gaits, but these gaits reflect that seen in the respective human disease.

Dopaminergic neurons preferentially degenerate when alpha-synuclein is panneuronally expressed, and their loss is implicated in the associated climbing defects^36^. We therefore asked whether other PD models that exhibit dopaminergic neuron degeneration show a similar gait profile. We examined a mutant of *parkin*, a conserved ubiquitin ligase that is one of the most common mutations underlying familial PD^39,43,44^ (Video 5). Interestingly, *park^1^* mutant flies also exhibited a decrease in hind leg domain length (Figs. 3Biv, 3Ciii, S5Ciii) and hind leg stride length compared to the mid legs (Figs. 3Biv, 3Ciii, S5Cvi). This resulted in lowered ratios of hind to mid leg domain length (Fig. 3Ciii, S5Civ) and stride length (Fig. 3Ciii, S5Cvii). Overall, the PD models *park^1^* and *Elav-Gal4>alpha-synuclein* showed gait profiles that were strikingly similar (Figs. 3Cii and 3Ciii), despite being genetically dissimilar. We wondered if hind leg rigidity was a basal property of poor motor function, and hence examined *mir-263a^KO^* flies, which also climb poorly^18^ (Fig. 3A). *Mir-263a^KO^* flies did not exhibit preferential hind leg rigidity (Fig. 3Civ, S5C), and its gait signature differed from that of SCA3 and PD flies (Figs. 3Ci-iv). The gait signature observed in alpha-synuclein and *park^1^* flies led us to hypothesise that dopaminergic neuron dysfunction may be the common underlying cause. To test this, we expressed mutant SCA3 under control of the dopaminergic driver ple-Gal4. Interestingly, while pan-neuronal expression of mutant SCA3 caused ataxic gait and hyperkinetic movements leading to enlarged mid and hind leg domains (Figs. 3Bi, 3Ci, S5A, Video 6), expression of mutant SCA3 with ple-Gal4 had a hypokinetic effect. While there was no change in the length of the mid leg domains, the hind leg domain length, and the hind/mid leg domain length ratios decreased (Figs. 3Cv, S5D). This led to a gait signature that more closely resembled that of *park^1^* and *Elav-Gal4>alpha-synuclein* flies (Fig. 3Cv vs 3Cii, iii). These data suggest that perturbation of dopaminergic neuron function in *Drosophila* leads to preferential rigidity of hind leg movements, and a specific gait profile.

**Figure 3.**
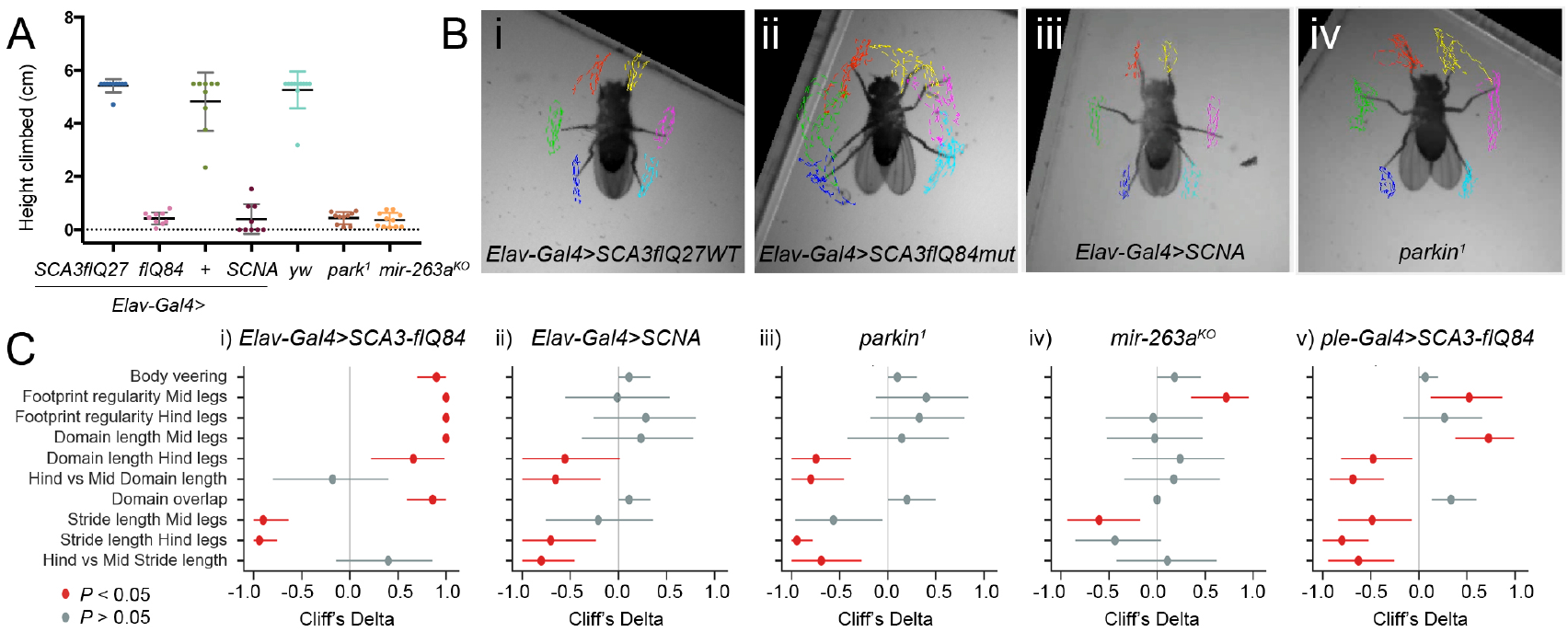
Gait signatures of *Drosophila* models of neurodegeneration reveal properties of underlying circuit dysfunctions. **A.** Climbing performance (highest height climbed in 30 s) of flies analysed. Data points are coloured as in Figure S5. **B.** Representative FLLIT-derived walking leg traces of the respective genotypes. **C.** Cliff’s delta indices of effect sizes (filled circles) of gait parameters with 95% confidence intervals (horizontal lines), with respective *P* values. Positive Cliff’s delta indicates an increase in mutant flies compared to respective controls, whilst negative Cliff’s delta indicates a decrease. Detailed statistics are given in Supplementary Table 1. Raw values are plotted in Figure S5. The following gait parameters were analysed: Body veering (Number of body turns normalised to the average number of strides per leg), Footprint regularity (Standard deviations of the anterior extreme position, normalised to body length), Leg domain lengths (normalised to body length), Average ratio of the hind vs mid domain length of the right and left sides, Domain overlap (number of pixels overlapping between leg domains, normalised to the average number of strides per leg), Stride lengths of the mid and hind legs (normalised to body length), Average ratio of the hind vs mid stride lengths of the right and left sides. \**P* < 0.05, \*\**P* < 0.01, \*\*\**P* < 0.001, \*\*\*\**P* < 0.0001. Genotypes examined: *Elav-Gal4>SCA3-flQ27* (n = 10), *Elav-Gal4>SCA3-flQ84* (n = 10), Elav-Gal4>+ (n = 9), *Elav-Gal4>SCNA* (n = 9), *yw* (n = 11), *park^1^* (n = 10), *mir-263a^KO^* (n = 11), *ple-Gal4>SCA3-flQ27* (n = 14), *ple-Gal4>SCA3-flQ84* (n = 15). *P* values were calculated using a non-parametric Mann-Whitney test except for *park^1^* and *mir-263a^KO^* which shared the same control (*yw*), hence, *P* was calculated using a non-parametric Kruskal-Wallis test with Dunn’s multiple comparisons *post-hoc* test (See Fig. S5). *See also Videos 2 - 6*.

## 6. Detection and characterisation of high frequency leg tremors in freely moving *Drosophila*

To determine if FLLIT can detect and quantify trembling leg movements in freely-walking flies, we first examined flies known to tremble: *Shaker* (*Sh^5^*) and *Hyperkinetic* (*Hk^2^*) mutants, which carry lesions in alpha and beta subunits of the Shaker voltage-gated potassium (Kv) channel, and exhibit leg shaking under ether anesthesia^45^. Aged, freely walking *Sh* mutants have also been described to exhibit “quivering” behaviour^17^, but the characteristics of these movements have not been previously quantified compared to wildtype flies.

Aged control (*yw*) flies exhibited infrequent sporadic shaking movements (Video 7), whereas *Hk^2^* mutants appeared to exhibit a more consistent tremor-like phenotype (Video 8). As such, wild-type legs followed relatively smooth paths, whereas leg paths of *Hk^2^* mutants were relatively irregular and uneven (Fig. 4A). To quantify these movements, local extrema of at least three pixels in amplitude were identified from the traces (circles and stars; Figs. 4A, B). A cutoff of three pixels was chosen to filter out small displacements that occur due to tracking errors (Fig. 2E), and roughly corresponded to the width of a leg tip (Fig. 2A).

*Hk^2^* mutants exhibited ~42 shaking events/s on average, whilst wild-type controls and *Sh^5^* mutants exhibited ~25 shaking events/s. However, this was not statistically significant (Fig. 4C). As tremors are periodic shaking movements, we then looked for tremor episodes: a series of three or more shaking events separated by < 100 ms (conservatively chosen based on the average stride rate of ~10 Hz in control flies in our assay; Fig. 4Ai). Each of these events was defined as a tremor event (red circles and stars; Figs. 4A, B). *Hk^2^* flies exhibited an average of ~8 tremor events per second, significantly more than control and *Sh^5^* flies, which showed almost no tremor events (*P* < 0.01)(Fig. 4D).

We determined the frequency of *Hk^2^* leg tremor episodes by examining the time interval between consecutive tremor events. A predominant inter-peak interval of 20–30 ms was observed (*P* < 0.01 by running a permutation test with 100,000 iterations; Fig. 4E), corresponding to a tremor frequency of ~33–50 Hz. Interestingly, *Hk^2^* leg tremors mostly occurred in the hind legs: An average of 6.7 tremor events per second were observed in hind legs of *Hk^2^* flies, accounting for ~89% of the tremors in each fly on average (Fig. 4F). Therefore, our method is able to detect and measure *Drosophila* leg tremors.

We then quantified tremor behaviour in the mutants that we previously examined for gait defects (Fig. 3). We found that *SCA3* mutants were the only flies to show tremor behaviour amongst the genotypes examined (Fig. 4G, Video 3). PD flies did not show show tremors when walking, nor did flies expressing SCA3 only in dopaminergic neurons (Fig. 4G). *Elav-G4>SCA3-flQ84* tremor showed a similar frequency to that of *Hk^2^* flies (*P* < 0.0001 by running a permutation test with 100,000 iterations; Fig. 4H). However, unlike *Hk^2^* flies, 95% of *SCA3* tremors occurred in the mid legs: An average of 9.9 tremor events per second were observed in mid legs of *Elav-G4>SCA3-flQ84*flies, accounting for ~97.5% of the tremors in each fly on average (Fig. 4Ii). A similar trend held when *Elav-G4>SCA3-flQ84* mutants flies walked upside-down/inverted– An average of 5.3 tremor events per second were observed in mid legs of inverted *Elav-G4>SCA3-flQ84* flies, accounting for ~95.7% of the tremors in each fly on average (Fig. 4Iii). This suggests that there are at least two different circuits whose perturbation can cause tremor.

**Figure 4.**
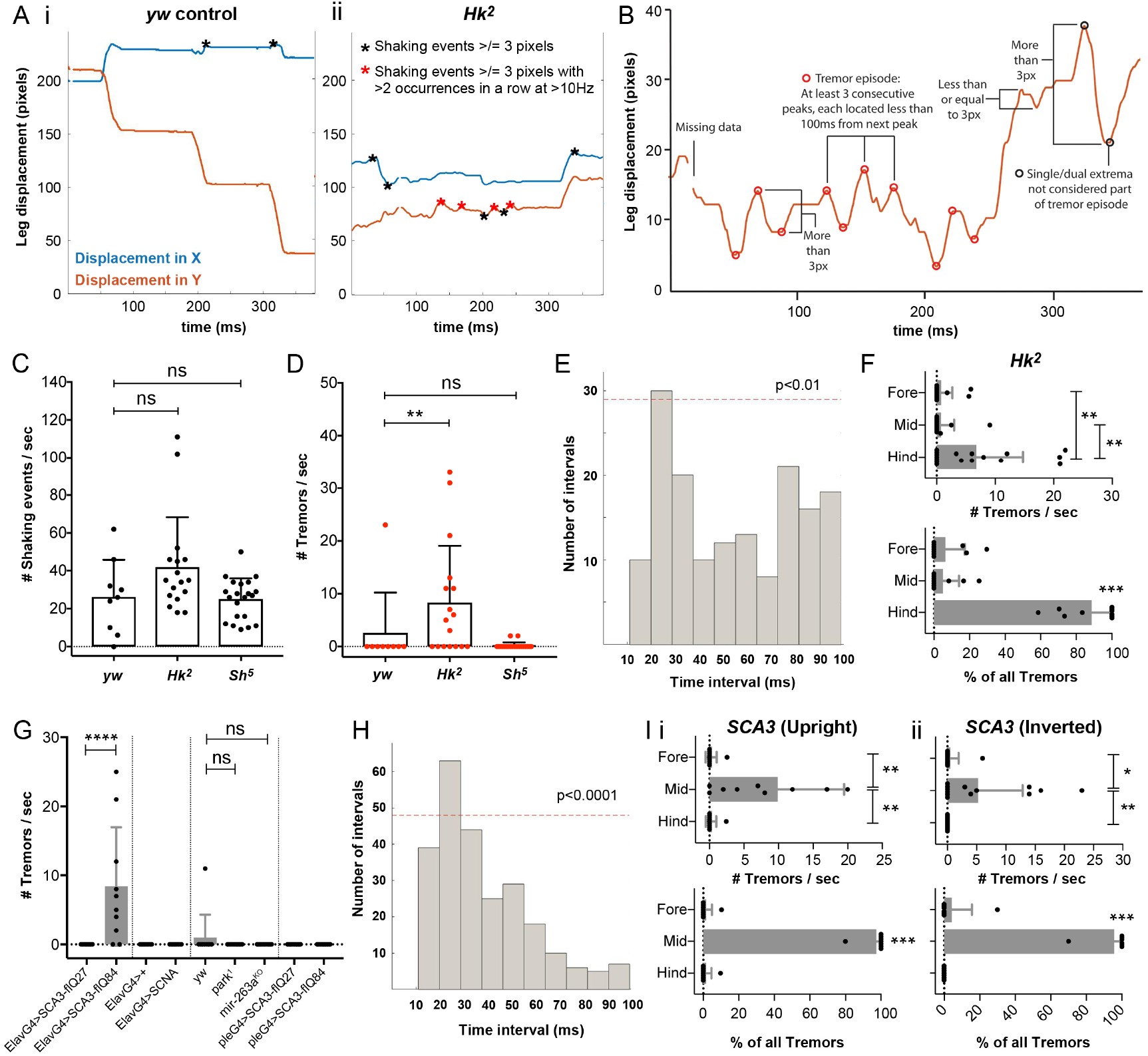
Detection and characterisation of high frequency leg tremors in *Drosophila* mutants show that at least two circuits underlie tremor. **A.** Representative leg traces of freely walking control (*yw*) and *Hk^2^* mutant *Drosophila*. Red indicates the displacement traces in the X direction, and blue indicates the displacement traces in the Y direction. Stars indicate the shaking events at least 3 pixels in size: black and red stars mark all shaking events, whereas red stars mark only tremor events occurring in three consecutive peaks or valleys (shown in B). **B.** Schematic of a representative trace showing the parameters used to determine shaking and tremor events. **C.** Number of shaking events in control (n = 11), *Hk^2^* (n = 17) and *Sh^5^* (n = 21) *Drosophila*. **D.** Number of tremor events in control, *Hk^2^* and *Sh^5^ Drosophila*. **E.** Distribution of the time interval durations between tremor peaks or valleys in *Hk^2^* flies. A significant proportion of events showed an interval duration of 20-30 ms (*P* < 0.01; *P* value was determined by running a non-parametric permutation test with 100,000 iterations), reflecting a tremor frequency of ~33-50Hz. **F.** Top: Number of tremors per second in the fore, mid and hind legs of each *Hk^2^* fly that exhibited tremors (n = 17, of which 10 flies showed a total of 140 tremors/s). Bottom: Percentage of all tremors accounted for by either the fore, mid or hind legs in each *Hk^2^* fly that exhibited tremors. **G.** Number of tremors per second exhibited by each of the genotypes examined: *Elav-Gal4>SCA3-flQ27* (n = 10), *Elav-Gal4>SCA3-flQ84* (n = 10), *Elav-Gal4>+* (n = 9), *Elav-Gal4>SCNA* (n = 9), *yw* (n = 11), *park^1^* (n = 10), *mir-263a^KO^* (n = 11), *ple-Gal4>SCA3-flQ27* (n = 14), *ple-Gal4>SCA3-flQ84* (n = 15). **H.** Distribution of the time interval durations between tremor peaks or valleys in Elav-Gal4>SCA3-flQ84 flies. A significant proportion of events showed an interval duration of 20-30 ms (*P* < 0.0001; *P* value was determined by running a non-parametric permutation test with 100,000 iterations), reflecting a tremor frequency of ~33-50Hz. **I.** Top: Number of tremors per second in the fore, mid and hind legs of each *Elav-Gal4>SCA3-flQ84* fly that exhibited tremors when (i) Walking upright (n = 10, of which 8 flies showed a total of 104 tremors/s), or (ii) Walking inverted (n = 15, of which 7 flies showed a total of 85 tremors/s). Bottom: Percentage of all tremors accounted for by either the fore, mid or hind legs in each *Elav-Gal4>SCA3-flQ84* fly that exhibited tremors when (i) Walking upright (n = 10), or (ii) Walking inverted (n = 15). *P < 0.05, **P < 0.01, ***P < 0.001, ****P < 0.0001. All data were analysed using a non-parametric Kruskal-Wallis test with Dunn’s multiple comparisons *post-hoc* test unless otherwise stated above. Bars represent the means and standard deviations. *See also Videos 7 and 8*.

## Discussion

Our study describes the development of a machine learning method that segments legs and tracks leg claw positions of freely moving flies captured on video, and its use to study gait and tremors in fly models of Parkinson’s Disease and Spinocerebellar ataxia 3. We demonstrate that machine learning can improve upon an approach that uses only morphological parameters, so that supervised learning is carried out without requiring the time-consuming manual annotation of training sets. Most of the FLLIT method is fully automatic; only for the correction of tracked data is user input required. The number of corrections required will depend on the types of gait defects in the genotypes of interest. Tracking using FLLIT allowed us to analyse almost 190,000 video frames in this study.

FLLIT does not require a contact surface for detection^19,20^ or leg markers for tracking^21^, is not sensitive to variations in recording parameters and is not rule-based^23^. These strengths permit its application to other animals, which we show by application to spiders. Images of segmented legs are saved by FLLIT, and could potentially be used for subsequent analyses. Tracked FLLIT data may be applied to other methods that use tracked and/or labelled data to identify and describe complex behaviours and patterns^46-48^, or combined with methods for circuit manipulation and functional imaging. While FLLIT outperforms state-of-the-art methods in our walking task, it is not optimal for all applications. The semi-automatic FlyLimbTracker^23^ would be more suitable if tracking of other leg segments is required. FlyWalker^19^ may be more suitable than FLLIT if only footprint information is required. Also, FLLIT requires a fixed viewing angle, from either above or below a walking fly. Recently developed deep learning approaches^25,26^ allow for tracking of varied behaviours and setups based on user-annotated training sets.

Using FLLIT, we characterise, for the first time, gait signatures of fly models of neurodegeneration. Interestingly, SCA3 and PD flies recapitulate distinct gait features of the respective human disease. SCA3 flies exhibit lurching, ataxic gait, while PD flies show stride rigidity, especially in the hind legs. These phenotypes are not differing in degree, rather, they are opposite in character: The former being hyperkinetic, and the latter, hypokinetic. Our study suggests that perturbation of dopaminergic circuits underlie the PD fly gait dysfunctions, resulting in a specific, striking gait signature in PD fly models that are otherwise genetically unrelated. Going forward, manipulation of other subsets of neurons may help us to understand how specific behavioural dysfunctions can result from perturbation of disease genes in different circuits. Classification of mutants with similar gait signatures may reveal novel relatedness of disease pathways and molecular mechanisms.

Tremor is increasingly prevalent in our aging population^49^, and improving our understanding of the mechanisms underlying tremor is important for the rational development of treatments. In this study, we detect and quantify leg tremor in freely-moving *Drosophila*, from large datasets of leg movement data tracked from high-speed video. To our knowledge, this is the first automated, image-based method for tremor measurement in any animal model. We propose that by combining these data with functional and imaging-based genetic tools, *Drosophila* models will be useful for understanding the mechanisms underlying tremor and tremor-associated diseases. Our analysis of *Hk^2^* and SCA3 mutants indicated that fly tremors occurred at 30–50 Hz. This is much more rapid than the tremors that occur in humans, such as those in Essential Tremor (4–12 Hz^50^), and orthostatic tremors (13–18Hz^51^). The reason for this can only be determined when the biophysical and cellular mechanisms underlying tremor are better understood. Our study only measured tremors that occured during walking, which are likely action tremors. Several other types of tremor exist in humans, including resting and postural tremors^52^. Notably, the PD flies examined did not exhibit action tremors. We also did not observe resting tremors in these flies; however, different assay conditions may need to be developed to systematically determine if other categories of tremor found in humans also occur in *Drosophila*. It will be now interesting to examine other mutants that exhibit tremors, to determine their different tremor “signatures”, and to compare these to corresponding human diseases.

In summary, we have developed an automated program for segmenting and determining leg-tip positions of freely-walking flies captured on high-speed video. Our method, FLLIT, uses machine learning, without the need for manual annotation of training sets. Using FLLIT to quantify gait and tremor characteristics of fly models of PD and SCA3, we find marked similarities in these models compared to the respective human disease. We identify a gait signature of PD flies and dopaminergic circuit dysfunction, and a signature of fly SCA3 tremors. These experiments set the stage for experimentation with more limited subsets of neurons in difference disease contexts, in order to understand how specific behavioural dysfunctions result from perturbation in different circuits, and for exploration of causal cellular and molecular pathways.

## Acknowledgements

The authors would like to thank Prof Paul Matsudaira (National University of Singapore) for helpful discussions, and Profs Stephen Cohen (University of Copenhagen) and Pavan Ramdya (École Polytechnique Fédérale de Lausanne) for comments on the manuscript. The authors would also like to thank Huilin Zhu, Chong Swee Tong, Jefferson Fu and Alice Liu for technical support and the Bloomington Drosophila Stock Centre (Indiana, USA) for making available the *Drosophila* strains used in this work. The authors would like to thank Dr. Jessica Tamanini of Insight Editing London for proofreading an earlier version of the manuscript. This work was supported by the Institute of Molecular and Cell Biology, Singapore; the Bioinformatics Institute, Singapore; the Agency for Science Technology and Research Joint Council Organization (grant number 15302FG149 to SA, LC, ACC and EKT and grant number 1431AFG120 to LC and ACC); and the Clinical Research Flagship Programme (Parkinson’s Disease) administered by the Singapore Ministry of Health’s National Medical Research Council (grant number NMRC/TCR/013-NNI/2014 to SA and EKT).

## Methods

### Iterative Training Module for Leg Segmentation

Leg segmentation is achieved using a supervised learning approach. Training images are automatically generated via image-processing steps, without user annotation.

First, a pool of representative images is obtained from a set of input images. In cases where the image set is an entire video, the representative images are obtained by uniform sampling from the video frames. To this aim, we select one image frame from every 20 frames. The operations described below are carried out for each image in the pool.

1. The silhouette of the subject animal (*Drosophila*) is extracted as a binary foreground via background subtraction, using the following formula: Silhouette =∣Image (x, y) − Background∣ > *Threshold*
2. Skeletonisation and edge detection of the silhouette foreground (standard image morphological operations) was performed on the images. The overlap between the skeleton image and the edge image primarily occur in the leg regions. The pixels within these regions are identified as positive samples containing the leg segments. The negative image consists of the fly body and background.
3. A fixed number of high-confidence samples are extracted after morphological operations on the segmented results, and used in a supervised learning approach. Each training sample is extracted as an image patch of 41 x 41 pixels and represented as an instance label pair (*x*_i_, *y_i_*), where *x_i_* denotes the image patch and *y*_i_ = ±1 denotes the corresponding label of the central pixel. This process provides the initial training dataset to learn the following Kernel-Boost classifier^53,54^:

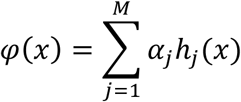

Where the classifier function φ (*x*) is a weighted sum of *M* = 100 weak learners *h_j_* with corresponding weights α*_j_* The primary framework of the Kernel-Boost classifier is gradient boosting, which adopts a greedy algorithm with quadratic approximation (Box 1).

#### Box 1

Greedy Algorithmwith Quadratic Approximation^55,56^

##### INPUT

Labelled training samples {(*x*_i_, *y*_i_)}

Exponential loss function *L*(*y*_i_, *φ*(*x*_i_)) = *e^yi^*φ(*x*i)

\# Iterations *M* = 100

1. Initialize model with: *φ*_0_(⋅) = 0
2. For *j*= 1 ∶ *M*
3. Compute the weight 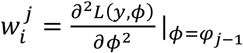 and the “pseudo-residual”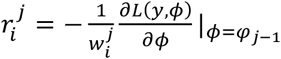
4. Update the *j*-th weak learner *h_j_*:

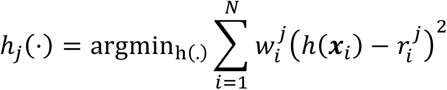
5. Update the j-th *α_j_* weight by line search:

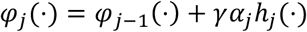
6. End for on j

##### OUPUT

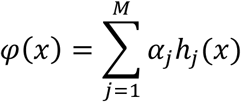

The Kernel-Boost classifier uses the following approach to update the weak learner of the algorithm (Box 1, step 4). The training set *T* is randomly split into two sets of fixed sizes:

> *T* = *T*1 ∪ *T*2

Instead of relying on pre-defined features, here features are automatically learnt on the first training set *T*1 in the form of convolution kernels. The weak learners are simultaneously learnt in the form of decision trees on the second training set *T*2. Essentially, the weak learners ℎ*_j_* will be a combination of kernels *K* and tree parameters *τ* (split threshold), *η* (leaf values).

### Learning Kernels on T1

The kernels are square windows, 4-19 pixels in length and operate on a specific fixed square region within each image patch *x_i_*. A total of 100 candidate kernels are obtained at boosting iteration *j*, with the p-th kernel **K***^j^_p_* being identified by:

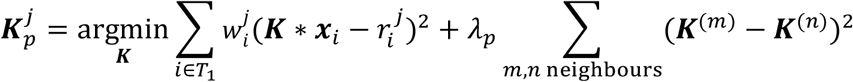

Where *K* ∗ *x_i_* denotes the convolution of kernel *K* on the fixed square region within image patch *x_i_*. The second term λ*_p_* Σ_m,n neighbours_ (*K*^(m)^ − *K*^(n)^)^2^ is a regularization term introduced to impose a smooth kernel. Here, *K*^(m)^ denotes the *m*-th pixel of kernel *K* and λ*_p_* Ʃ_*m,n*,neighbours_(**K**^*(m)*^ – **K**^*(n)*^)^2^ is a regularization factor for the p-th kernel that is randomly assigned to one of three values: 100, 500 or 1,000.

### Constructing Decision Tree on T2

Decision tree learning is performed one split at a time, up to a depth of five levels (32 leaf nodes). To learn the tree parameters (split threshold τ*^j^* and leaf values t a decision node, a split search is first performed on T2 for every candidate kernel *K_p_^j^* as follows:

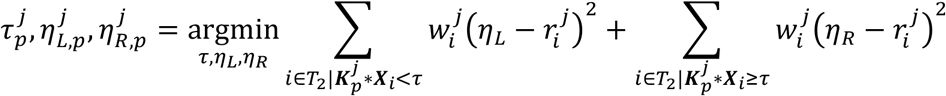

Where τ^j^_p_ is the split threshold and η*^j^_L,p_*, *^j^_R,p_* are the leaf values for a specific kernel *K_p_^j^*. In actual implementation, the optimal split threshold *τ_p_* is found by exhaustive search. Subsequently, the leaf values η*^j^_p_* are simply given by the weighted sum:

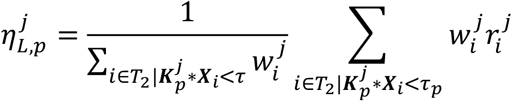

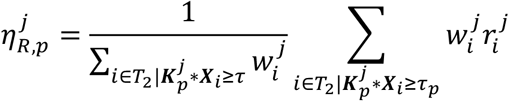

The split cost is evaluated for each candidate kernel *K_p_^j^*:

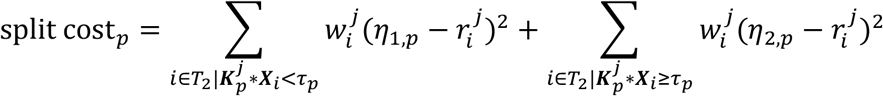

The kernel giving the smallest split cost is chosen for this split.

Subsequently, the split search is performed recursively up to the desired tree depth and the final output for the *j*-th weak learner is the set of kernels *K_p_^j^* as well as the decision tree. A summary of the parameters used is shown in Table S1.

### Prediction with the Kernel-Boost classifier

During the prediction phase, the classifier is applied onto the subject animal (*Drosophila*) silhouette foreground. For each pixel, the learned kernels and decision tree splitting are applied onto the image patches. Passing through all iterations gives a confidence probability that the pixel of interest belongs to a leg. From our experiments, we found that a confidence threshold of 0.6–0.65 was optimal.

**Table S1:** Parameters used in the Kernel-Boost Algorithm

| Parameter | Value |
| --- | --- |
| Number of Positive Samples | 30000 |
| Number of Negative Samples | 30000 |
| Sample image patch size | 41 square pixels |
| Number of samples for learning convolutional kernels | 10000 |
| Min. Kernel Size | 4 square pixels |
| Max. Kernel Size | 19 square pixels |
| Number of kernels to explore for training one weak learner | 100 |
| Regularization factor | 100;500;1000 |
| Max. Decision Tree Depth | 5 |
| Shrinkage Factor $\gamma$ | 0.1 |

### Tracking module

For tracking, the segmented leg pixels are promoted from pixel level to object level representation by grouping connected sets of identified leg pixels while rectifying irregularities and noise arising during segmentation.

By obtaining the centroid and angle-of-rotation from the silhouette image, each frame is translated and rotated such that the subject animal (*Drosophila*) is aligned with the y-axis. After a single pixel-wide skeletonisation of the legs, the leg claw is identified as the endpoint at a maximum distance from the fly body. Tracking is automatically initialized by identifying the first frame with the correct number of identified legs, which are then labelled according to their geometric positions. In the case of *Myrmaplata plataleoides* (salticid spider) spider-leg tracking was manually initiated by marking each leg tip in the first frame. Subsequent tracking proceeds as normal.

Leg tracking invokes an optimal linear assignment problem where the leg tips in the next frame have to be consistently labelled as those in the current frame, as follows:

Given tips *x_i_ ^j^*the position of the *j*-th leg in frame *i* (in body centred coordinates), consistent labels *j* = 1,…, 6 have to be assigned to *x*_i+1_ of frame *i* + 1.

The assignment cost is the distance between the tips across the frames and the problem seeks a global minimization of this distance to assign a label to the tips in the next frame. Formally, the problem can be stated as:

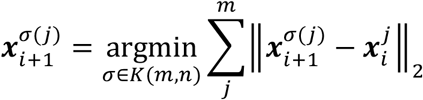

Where there are *m* tips identified in frame *i* and *n* tips identified in frame *i* + 1 and *σ* is an injective mapping from the set of *m* elements to the set of *n* elements, subject to the constraint that 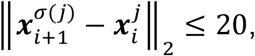, (i.e. the tip of a leg cannot move >20 pixels across a single frame).

We denote the set of all such possible mappings as K(*m*, *n*), using the Hungarian method^55,56^ *O*(*m*^3^) combinatorial algorithm. In the event that legs are not found, the last located position of the missing legs in a previous frame is utilised to match with any left-over tips in frame *i*+1. Similarly, in cases of leg occlusion or false negatives in identifying leg tips, the previous frame’s history is used to restore the correct identity upon leg tip re-appearance. A summary of the tracking procedure and parameters is shown in Box 2. A manual correction feature in the FLLIT program allows the user to correct mislabelled legs or make adjustments to tip positions.

#### Box 2

Tracking Procedure

##### INPUT

Segmented leg pixels in each frame

Drosophila Silhouette in each frame

Maximum Distance Constraint moved by a leg tip

1. For *i* = 1 ∶ Total number of frames
2. Obtain centroid position and orientation of the *Drosophila* from the binary silhouette. Translation and rotation operation to align the drosophila along the y-axis.
3. Group segmented leg pixels as different legs and identify leg tips as the endpoints at a maximum distance from the body. This gives all leg tips *x_i_* across all frames.
4. End for *i*
5. Tracking initialization: identify the starting frame with all legs being visible, and label them according to geometric position. This gives *x^j^_start_* for *j* = 1,…, #legs.
6. For *i* = Starting Frame + 1 ∶ End Frame
7. Labelling tips in frame 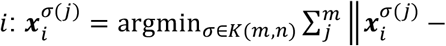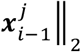 subject to the constraint 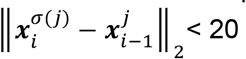 pixels with Hungarian algorithm.
8. Recovering missing tips with last seen information and left-over tips:

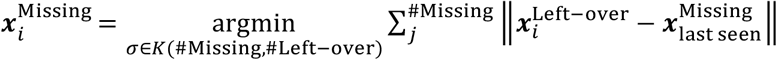
9. End For

##### OUTPUT

Trajectory of leg tips from start to end frame in both the body-centred as well as arena-centred frames of reference

## Experimental Methods

### System setup and video recording

Figures 1 and S1 illustrate the video setup. In brief, a Photron FastCam MC2 high speed camera was mounted below an arena containing the sample. The arena was backlit with a diffused infrared LED array. For *Drosophila*, the floor and ceiling of the arena consisted of glass microscope slides and the arena walls were cut from 1.5 mm thick transparent acrylic sheets, representing the height of the arena. For *Myrmaplata plataleoides* video recordings, the floor, ceiling and walls of the arena were cut from transparent acrylic sheets. The walls of the arena were 6.5 mm high.

*Drosophila* were transferred into the arena by mouth aspiration or after brief incapacitation by cooling on ice, and allowed to acclimatize for at least 5 min (mouth aspiration) or 15 min (on ice) before starting the recording. The field of view ranged from 9 mm x 9 mm square, to 12 mm x 12 mm square (10 mm x 10 mm default unless otherwise indicated). For *M. plataleoides*, the field of view measured 50 mm x 50 mm. Video recordings were carried out at 1,000 frames per second, with 512 x 512 pixel resolution at 25°C, unless otherwise indicated. Videos were captured when the animal walked straight through the middle of the arena, without touching the perimeter. For efficient automated background subtraction by FLLIT, only videos where the subject animal moved at least 1.5 body lengths were used. If the user wishes to analyse videos where the subject did not traverse at least 1.5 body lengths, they should separately upload a background image (see below).

### Animal handling

*Drosophila* stocks (*Sh^5^* (BL111), *Hk^2^* (BL55), *Elav-Gal4* (BL8765), *ple-Gal4* (BL8848), *UAS-SCNA* (BL51376) *SCA3-flQ27* (BL33609), *SCA3-flQ84* (BL33610) and *park^1^* (BL34747)) were obtained from the Bloomington Drosophila Stock Centre (Indiana, USA). *Mir-263aGal4^KO^/bft^24^* flies (referred to as *mir-263a^KO^*) were previously described^18^. Flies were reared at 25 ± 1°C in 70% relative humidity in an environmentally controlled incubator on a 12 h light-dark schedule. Crosses were set up with 20 females per bottle and flipped every 2 days to prevent overcrowding. Groups of 15–20 males were collected within 24 hr of eclosion and aged without further CO2 exposure. Flies were flipped onto fresh food every 2–3 days, and vials were laid on their sides to minimise flies getting stuck in the food. For ground truth, wildtype flies were analysed at 4-7 days. For the mutant genotypes, the age chosen for analysis was one at which ~50% of flies climbed below 1.5 cm (4^th^ etching on tubes used; see single fly climbing assay protocol below); and flies that climbed between 0.3-0.9cm were used for recording (2^nd^ and 3^rd^ etchings on tubes used). Based on this criteria, *yw*, *Sh^5^ and Hk^2^* aged flies (Fig. 4) were analysed at 35-41 days. *Elav-G4>SCA3-flQ27* and *Elav-G4>SCA3-flQ84* flies were analysed at 20-25 days. *Elav-G4>+* and *Ela-vG4>SCNA* flies were analysed at 48 days. *Yw* and *park^1^* flies were analysed at 35 days, *mir-263a^KO^* flies were analysed at 22-24 days. *Ple-G4>SCA3-flQ27* and *ple>SCA3-flQ84* flies were analysed at 21-25 days. All data provided are from males walking upright, except for *yw*, *Sh^5^ and Hk^2^* aged flies in Fig. 4, which were recorded walking upside down/inverted.

*M. plataleoides* were collected in Singapore and housed individually in cylindrical cages (6.5 cm x 8.5 cm) and reared to maturity in captivity. They were reared at 25 ± 1 °C and 80-90% relative humidity, on a 12 h light-dark schedule. The spiders were fed six fruit flies (*D. melanogaster*) twice a week with access to water *ad libitum*.

### Segmentation groundtruth

To empirically determine the classification threshold for leg pixel segmentation, we manually identified leg pixels in image frames randomly sampled from videos of wild-type *Drosophila*, taken using default recording parameters (8 videos, 2–3 images per video). We then determined the precision, recall and F_0.5_ scores achieved using the FLLIT segmentation classifiers.

**Figure M1:**
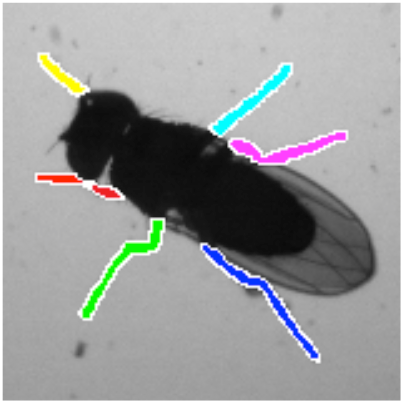
Representative image of an annotated, wild-type (*yw*) *Drosophila*.

The blurry regions on the edges of the legs are marked with white pixels and given a weight of 0, so that they will not be important in assessing the precision and recall scores (Fig. M1). The F_0.5_ score for the tested classifiers peaked at 0.6–0.65 (Fig. S2A); hence, we selected this threshold for subsequent analyses. Users may adjust the classification threshold in the FLLIT interface based on their requirements.

### Tracking groundtruth

Users were instructed to zoom in to the image frame to 800% magnification, and to label the leg-tip pixel as accurately as possible with no time constraint. One in every 20 frames (5% of each video) was annotated in this way. Different colours were used to label each leg tip, and labelled positions were then compared with leg-tip positions that were derived computationally or from another user.

### Error identification

Misidentification errors occurred mainly during leg retraction and occlusion, and had to be corrected to prevent error propagation. Corrections of these errors were carried out to minimize the need for further corrections, while allowing for missing data. We took advantage of the algorithm’s ability to match tips only within a set distance threshold (20 pixels) across frames, by locating a correction <20 pixels away from where the leg may reappear after occlusion, and >20 pixels away from the site of misidentification, to prevent subsequent further cases of misidentification (Fig. S2C). To avoid using these incorrectly annotated positions, sharp movements occurring within 1 ms were filtered out before carrying out subsequent analyses. Tips that were reported absent as a result of these corrections were included in the missing data tally.

### Background loading

Segmentation and tracking accuracy depend on clean background subtraction (the first step of image processing). As such, an automated background subtraction step was built into FLLIT. This automated background subtraction algorithm requires the subject animal to move at least a distance of 1.5 body lengths; hence, videos were made with this criteria in mind. All data shown were generated using the FLLIT-derived automated background subtraction. In most cases, this procedure performed well; in some cases, or if the subject animal does not traverse at least 1.5 body lengths, loading of a background can substantially improve segmentation and tracking (Fig. S2D). A manual background can be made either by taking a separate image of the background alone, or by constructing one via image processing.

## Side by side comparison to DeepLabCut^26^

### Dataset Preparation

Two manually-annotated *Drosophila* datasets with default settings were used as the ground truth datasets. Within each dataset, 200 training images were randomly selected and the training set was prepared according to the DeepLabCut specifications. A grayscale image was converted to three image channels by repeating the grayscale image channel. Each training set consisted of 200 three-channel images together with the respective annotated tip positions of the six legs. The remaining images within the two ground truth datasets formed the testing set.

### Training Parameters

The default training hyperparameters in DeepLabCut were used. A pre-trained ResNet 50 network was used as the convolutional layers for extracting image features. The Huber loss was adopted as the training loss with location refinement set to true. Training was done for 100,000 iterations separately on each training set.

### Testing phase

The model trained on one dataset was tested on both the test sets. We observed that a model trained with images from dataset A will often mismatch the leg identities on the test images from dataset B and vice versa. We hence manually corrected the leg identities to improve the results from DeepLabCut. These leg-tip positions labelled by DeepLabCut were then compared to the manually annotated dataset to generate the deviation in tip position in pixels.

## Side by side comparison to Isakov et. al.^22^

The above two ground truth datasets that were used for comparison to DeepLabCut were used. As the TDM method was sensitive to image contrast, each video was manually reviewed to select a section with the longest bout of straight walking whose image contrast was acceptable by the method for successful tracking. The tracking algorithm in Isakov et al. was then used to generate predictions for the fly centroid and tip positions of the legs. These leg-tip predictions labeled by Isakov et. al.^22^ were then compared to the manually annotated dataset to generate the deviation in tip position in pixels.

## Gait parameters

The following gait parameters were analysed: Body veering (Number of body turns normalised to the average number of strides per leg), Footprint regularity (Standard deviation of the anterior extreme position, normalised to body length), Leg domain length normalised to body length, Average ratio of the hind vs mid domain length of the right and left sides, Number of pixels overlapping between leg domains, normalised to the average number of strides per leg), Stride lengths of the mid and hind legs normalised to body length, Average ratio of the hind vs mid stride lengths of the right and left sides.

## Analysis of shaking and tremor events

Shaking and tremor events were analyzed as described (Fig. 4B). Matlab scripts for detecting extrema and determining tremor frequency are provided at the project website. For reference, the three tremor events highlighted in the tremor episode in Fig. 4B occurred over a period of ~80 ms, or ~37Hz (Video 3; at interval ~1450–1550 ms).

## Data preparation and handling

Videos were cropped so that the animal did not touch the perimeter of the arena during the trial. These videos were converted to TIFF format and analysed with FLLIT. Individual video and tracking data that support the findings of this study are available from the corresponding author upon reasonable request.

## Statistical analyses

Since several distributions did not conform to normality, we used non-parametric methods for statistical analyses. For comparing two genotypes, we used the Mann-Whitney test. For comparing three or more genotypes, we used the Kruskal-Wallis test with Dunn’s multiple comparisons *post-hoc* test. Statistical analysis was carried out using Prism 6 (GraphPad Software).

For gait signature analysis, we computed Cliff’s delta for 10 different gait parameters (Table 2). Cliff’s delta is a nonparametric measurement of effect size that reflects the likelihood that an observation from a test group is greater than an observation from a control group. It ranges from −1 (when all values in the mutant group are smaller than the control) to +1 (when all values in the mutant group are larger than the control). The greater the overlap between the two distributions, the closer to 0 Cliff’s delta will be. Unlike Cohen’s *d*, Cliff’s delta can be applied on non-normal distributions. Each mutant genotype was compared to the appropriate control as shown in Figure S5. Cliff’s delta was calculated according to the original formulation by Norman Cliff^57^ using NumPy <https://www.numpy.org>. 95% confidence intervals (95CIs) were obtained via bootstrap methods^58^ with 10,000 samples, using the scikits.bootstrap package <https://github.com/cgevans/scikits-bootstrap>. Forest plots were created with matplotlib <https://matplotlib.org/>.

## Body size measurement

Three still images from the video of each fly (first, middle and last frame) were used for measurement. Using Microsoft Paint, each image frame was magnified to 800%, and the anterior-most pixel of the head and posterior-most pixel of the abdomen at the midline were labelled. The labelled images were then opened in imageJ, and the scale was input accordingly (Distance in pixels: 512, Known distance: varies, Unit of length: mm). A line was drawn between the labelled head and abdomen tip pixels to obtain the body length. The length determined in each of the three images was then averaged to obtain the average body size.

## Single fly climbing assays

Climbing experiments were carried out between 3 and 6 pm to minimize circadian differences. Single flies were transferred to 14ml falcon tubes (Falcon #352059), with cut ends sealed with transparent plastic, and allowed to acclimate undisturbed for 15–30 min before testing. Flies were lightly tapped down to the bottom of the tube, and the climbing height attained in 30 s was measured. Tubes were then placed horizontally and retested 10–15 mins later. The average of the 2 technical replicates for each vial was recorded, and the height climbed for each fly was plotted as a single point.

## FLLIT software

The most recent version of FLLIT can be downloaded from https://github.com/BII-wushuang/FLLIT.

## User workflow

Users first select a data folder of image frames for analysis. They have an option to load a background, which is only necessary if the subject moves slowly or not at all through a portion of the video, as this hampers automated background subtraction. Training then automatically occurs on a subset of the dataset before segmentation is carried out. The confidence threshold for classification can also be selected. Default settings are pre-loaded. Segmentation then proceeds without the need for further input. After segmentation, tracking begins. Leg claw positions are highlighted and labelled during tracking. Errors in leg identity assignment may occasionally occur when legs come in close proximity to each other, cross over, or when a leg emerges after a prolonged period of time being hidden. As leg claw positions are tracked across adjacent frames, correcting a mis-labelled leg identity at the earliest point following the wrongful assignment is usually sufficient to correct leg identity in subsequent frames. The user can perform error correction either during tracking, or after tracking is completed. Tracking can then be resumed from the point of correction.

